# Low neutralization of SARS-CoV-2 Omicron BA.2.75.2, BQ.1.1, and XBB.1 by 4 doses of parental mRNA vaccine or a BA.5-bivalent booster

**DOI:** 10.1101/2022.10.31.514580

**Authors:** Chaitanya Kurhade, Jing Zou, Hongjie Xia, Mingru Liu, Hope C. Chang, Ping Ren, Xuping Xie, Pei-Yong Shi

## Abstract

The newly emerged SARS-CoV-2 Omicron BQ.1.1, XBB.1, and other sublineages have accumulated additional spike mutations that may affect vaccine effectiveness. Here we report neutralizing activities of three human serum panels collected from individuals 1-3 months after dose 4 of parental mRNA vaccine (post-dose-4), 1 month after a BA.5-bivalent-booster (BA.5-bivalent-booster), or 1 month after a BA.5-bivalent-booster with previous SARS-CoV-2 infection (BA.5-bivalent-booster-infection). Post-dose-4 sera neutralized USA-WA1/2020, BA.5, BF.7, BA.4.6, BA.2.75.2, BQ.1.1, and XBB.1 SARS-CoV-2 with geometric mean titers (GMTs) of 1533, 95, 69, 62, 26, 22, and 15, respectively; BA.5-bivalent-booster sera improved the GMTs to 3620, 298, 305, 183, 98, 73, and 35; BA.5-bivalent-booster-infection sera further increased the GMTs to 5776, 1558,1223, 744, 367, 267, and 103. Thus, although BA.5-bivalent-booster elicits better neutralization than parental vaccine, it does not produce robust neutralization against the newly emerged Omicron BA.2.75.2, BQ.1.1, and XBB.1. Previous infection enhances the magnitude and breadth of BA.5-bivalent-booster-elicited neutralization.

## Main

The continuous emergence of new variants of severe acute respiratory syndrome coronavirus 2 (SARS-CoV-2) has caused global waves of infection. Since its first report in November 2021 in South Africa, Omicron has become the dominating variant due to its high transmissibility and immune evasion [1, 2]. Many Omicron sublineages have emerged over time. The initial Omicron BA.1 was displaced by BA.2, which has further evolved to sublineages BA.2.12.1, BA.2.75, BA.2.75.2, BA.4, and BA.5, among which BA.5 is currently dominant in many parts of the world. BA.4 and BA.5 have an identical spike sequence (defined as BA.4/5 hereafter) and their offsprings BA.4.6, BF.7, and BQ.1.1 are expanding prevalence in circulation. As of October 29, 2022, the BA.2-derived sublineage BA.2.75.2 accounted for 1.8% of the total SARS-CoV-2 infection in the United States; whereas the BA.4/5-derived sublineages BA.4.6, BF.7, BQ.1, and BQ.1.1 accounted for 9.6%, 7.5%, 14%, and 13.1% of the total infection, respectively; all these sublineages were on uptick trajectory last month (https://covid.cdc.gov/covid-data-tracker/#variant-proportions). In addition, the Omicron sublineage XBB, first identified in India in August 2022, is rapidly spreading in Europe and has been detected in the United States. XBB was predominant in Singapore, accounting for 54% of local infections during October 3-9, 2022 (Ministry of Health, Singapore-https://www.moh.gov.sg/).

SARS-CoV-2 spike mutations often contribute to immune evasion and/or transmission efficiency [3–9]. Previous studies showed that 3 or 4 doses of parental mRNA vaccine did not elicit robust neutralization against BA.4/5, supporting the bivalent vaccine strategy [10–12]. Since the newly emerged Omicron sublineages have accumulated additional spike mutations (**Fig. 1A**), it is important to examine the vaccine-elicited neutralization against these new sublineages. The goal of this study was to compare the neutralizing activities against six newly emerged Omicron sublineages (BA.5, BF.7, BA.4.6, BA.2.75.2, BQ.1.1, and XBB.1) using human sera collected from individuals who received 4 doses of parental mRNA vaccine or a BA.5-bivalent-booster.

**Figure 1.**
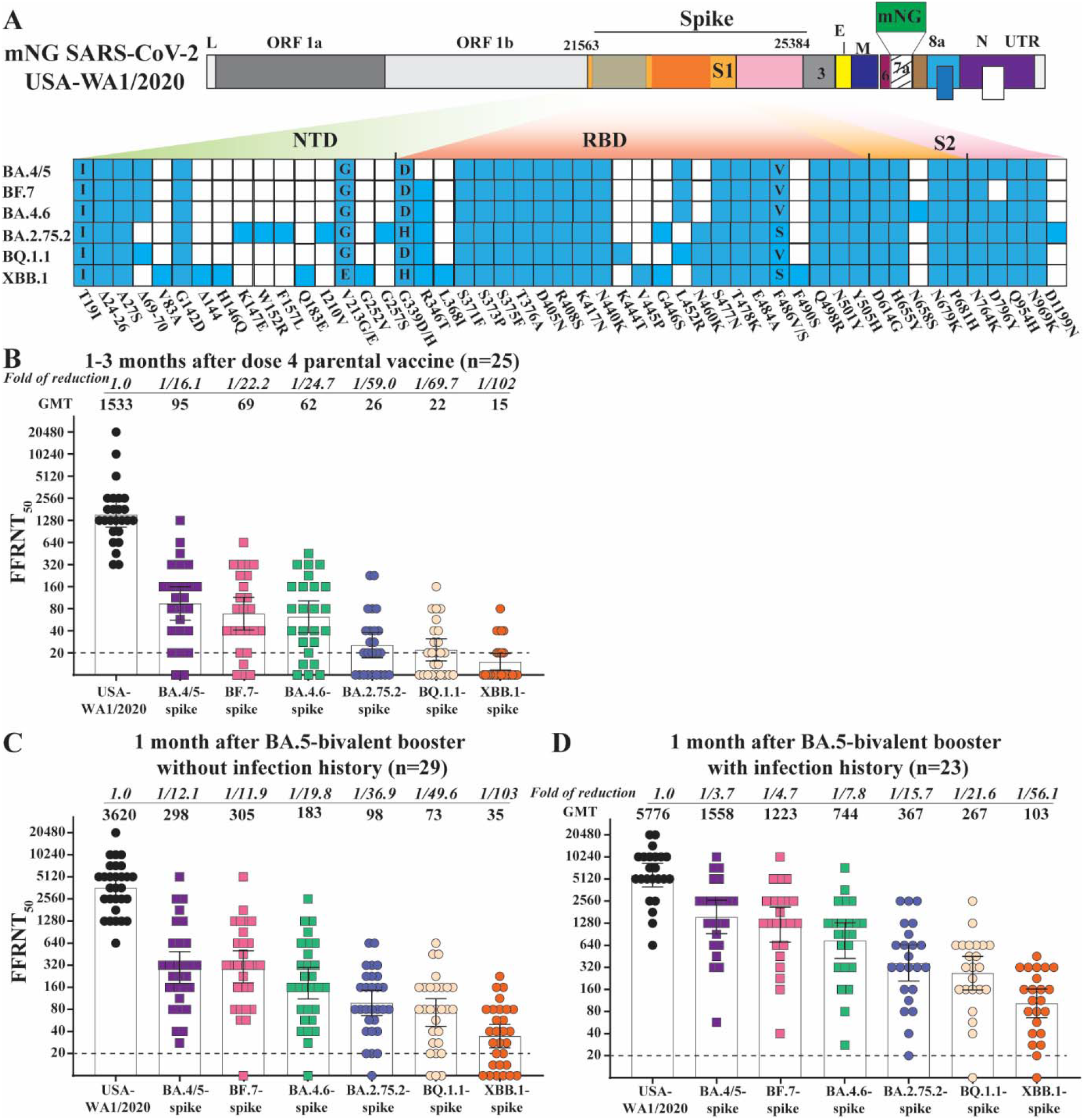
Neutralization of Omicron sublineages after 4 doses of parental mRNA vaccine or a BA.5-bivalent-booster. (A) Construction of Omicron sublineage-spike mNG SARS-CoV-2. mNG USA-WA1/2020 SARS-CoV-2 was used to engineer Omicron-spike SARS-CoV-2s. The mNG reporter gene was engineered at the open-reading-frame-7 (ORF7) of the USA-WA1/2020 genome. Amino acid mutations, deletions (Δ), and insertions (Ins) are indicated for variant spikes in reference to the USA-WA1/2020 spike. L: leader sequence; ORF: open reading frame; NTD: N-terminal domain of S1; RBD: receptor binding domain of S1; S: spike glycoprotein; S1: N-terminal furin cleavage fragment of S; S2: C-terminal furin cleavage fragment of S; E: envelope protein; M: membrane protein; N: nucleoprotein; UTR: untranslated region. (B) FFRNT_50_s of human sera after dose 4 parental mRNA vaccine. Human sera were collected 1-3 months after dose 4 parental vaccine. The bar heights and the numbers above indicate neutralizing GMTs. The whiskers indicate 95% CI. The fold of GMT reduction against each Omicron sublineage, compared with the GMT against USA-WA1/2020, is shown in italic font. The dotted line indicates the limit of detection of FFRNT_50_. FFRNT_50_ of <20 was treated as 10 for plot purposes and statistical analysis. The *p* values (Wilcoxon matched-pairs signed-rank test) for group comparison of GMTs are the following. USA-WA1/2020 versus all Omicron sublineage-spike: <0.0001; BA.4/5-spike versus BF.7-, BA.4.6-, BA.2.75.2-, BQ.1.1-, XBB.1-spike: 0.029, 0.001, <0.0001, <0.0001, and <0.0001, respectively; BF.7-spike versus BA.4.6-, BA.2.75.2-, BQ.1.1-, XBB.1-spike: 0.103, <0.0001, <0.0001, and <0.0001, respectively; BA.4.6-spike versus BA.2.75.2-, BQ.1.1-, XBB.1-spike: 0.0001, <0.0001, and <0.0001, respectively; BA.2.75.2-spike versus BQ.1.1-, XBB.1-spike: 0.24 and <0.0001, respectively; BQ.1.1-spike versus XBB.1-spike: 0.0028. (C) FFRNT_50_ of 29 sera collected 1 month after a BA.5-bivalent-booster from individuals without infection history. The *p* values (Wilcoxon matched-pairs signed-rank test) for group comparison of GMTs are the following. USA-WA1/2020 versus all Omicron sublineage-spike: <0.0001; BA.4/5-spike versus BF.7-, BA.4.6-, BA.2.75.2-, BQ.1.1-, XBB.1-spike: 0.844, <0.0001, <0.0001, <0.0001, and <0.0001, respectively; BF.7-spike versus BA.4.6-, BA.2.75.2-, BQ.1.1-, XBB.1-spike: all <0.0001; BA.4.6-spike versus BA.2.75.2-, BQ.1.1-, XBB.1-spike: all <0.0001; BA.2.75.2-spike versus BQ.1.1-, XBB.1-spike: 0.69 and <0.0001, respectively; BQ.1.1-spike versus XBB.1-spike: <0.0001. (D) FFRNT_50_ of 23 sera collected 1 month after BA.5-bivalent-booster from individuals with infection history. The *p* values (Wilcoxon matched-pairs signed-rank test) for group comparison of GMTs are the following. USA-WA1/2020 versus all Omicron sublineage-spike: <0.0001; BA.4/5-spike versus BF.7-, BA.4.6-, BA.2.75.2-, BQ.1.1-, XBB.1-spike: 0.0049, <0.0001, <0.0001, <0.0001, and <0.0001, respectively; BF.7-spike versus BA.4.6-, BA.2.75.2-, BQ.1.1-, XBB.1-spike: all <0.0001; BA.4.6-spike versus BA.2.75.2-, BQ.1.1-, XBB.1-spike: 0.0005, <0.0001, and <0.0001, respectively; BA.2.75.2-spike versus BQ.1.1-, XBB.1-spike: 0.114 and <0.0001, respectively; BQ.1.1-spike versus XBB.1-spike: <0.0001.

To facilitate neutralization measurement, we engineered the complete spike gene from Omicron sublineages BA.4/5 (BA.4: GISAID EPI_ISL_11 542270; BA.5: GISAID EPI_ISL_11542604), BF.7 (GISAID EPI_ISL_14425795), BA.4.6 (GISAID EPI_ISL_15380489), BA.2.75.2 (GISAID EPI_ISL_14458978), BQ.1.1 (GISAID EPI_ISL_15542649), or XBB.1 (GISAID EPI_ISL_15232105) into the backbone of mNeonGreen (mNG) reporter USA-WA1/2020 SARS-CoV-2 [13], a strain isolated in January 2020 (**Fig. 1A**). Compared with wildtype USA-WA1/2020, insertion of mNG gene at open-reading-frame-7 of the viral genome attenuated the virus *in vivo* [14]. So, recombinant mNG USA-WA1/2020 bearing variant-spike can be safely used for authentic SARS-CoV-2 neutralization and antiviral screening [15]. Passage 1 of recombinant BA.4/5-, BF.7-, BA.4.6-, BA.2.75.2-, BQ.1.1-, and XBB.1-spike mNG USA-WA1/2020 viruses were sequenced to ensure no undesired mutations. Only the passage 1 virus stocks were used to determine the 50% fluorescent focus-reduction neutralization titers (FFRNT_50_) of vaccinated human sera.

Three human serum panels with distinct vaccination and/or SARS-CoV-2 infection history were analyzed. The first panel consisted of 25 sera obtained from individuals 1-3 months post dose 4 of Pfizer/BioNTech or Moderna parental mRNA vaccine (post-dose-4 sera). The second panel consisted of 29 sera collected from individuals 1 month post BA.5-bivalent-booster (BA.5-bivalent-booster sera). All sera from the first and second panels were tested negative against viral nucleocapsid protein, suggesting no previous SARS-CoV-2 infection. The third panel consisted of 23 sera collected from individuals who were previously infected by SARS-CoV-2 (nucleocapsid antibody positive) and received a BA.5-bivalent-booster 1 month ago (BA.5-bivalent-booster-infection sera). All participants from the second and third panels had also received 2, 3, or 4 doses of parental mRNA vaccine before receiving the BA.5-bivalent-booster. **Tables S1-3** summarize the serum information and neutralization for each serum panel.

Post-dose-4 sera neutralized USA-WA1/2020-, BA.4/5-, BF.7-, BA.4.6-, BA.2.75.2-, BQ.1.1-, and XBB.1-spike mNG SARS-CoV-2 with geometric mean titers (GMTs) of 1533, 95, 69, 62, 26, 22, and 15, respectively (**Figure 1B** and **Table S1**). The neutralizing GMTs against BA.4/5-, BF.7-, BA.4.6-, BA.2.75.2-, BQ.1.1-, and XBB.1-spike viruses were 16.1-, 22.2-, 24.7-, 59-, 69.7-, and 102-fold lower than the GMT against the USA-WA1/2020-spike virus, respectively (**Figure 1B**). Compared with the GMT against the current dominant BA.4/5, the neutralizing GMTs against BF.7-, BA.4.6-, BA.2.75.2-, BQ.1.1-, and XBB.1-spike viruses were reduced by 1.4-, 1.5-, 3.7-, 4.3-, and 6.3-fold, respectively. The GMTs against BA.2.75.2 (26) and BQ.1.1 (22) were barely above 20, the detection limit of FFRNT; whereas the GMT against XBB.1 (15) was below the FFRNT detection limit. These results indicate that (i) 4 doses of parental mRNA vaccine do not elicit robust neutralization against the newly emerged Omicron sublineages and (ii) the rank of neutralization evasion is in the order of BA.4/5 < BF.7 ≤ BA.4.6 < BA.2.75.2 ≤ BQ.1.1 < XBB.1.

BA.5-bivalent-booster sera neutralized USA-WA1/2020-, BA.4/5-, BF.7-, BA.4.6-, BA.2.75.2-, BQ.1.1-, and XBB.1-spike SARS-CoV-2s with GMTs of 3620, 298, 305, 183, 98, 73, and 35, respectively (**Figure 1C** and **Table S2**). The neutralizing GMTs against BA.4/5-, BF.7-, BA.4.6-, BA.2.75.2-, BQ.1.1-, and XBB.1-spike viruses were 12.1-, 11.9-, 19.8-, 36.9-, 49.6-, and 103-fold lower than the GMT against the USA-WA1/2020, respectively (**Figure 1C**). The data indicate that (i) BA.5-bivalent-booster elicits significantly higher neutralizing titers against the recently emerged Omicron sublineages than 4 doses of parental vaccine and (ii) despite this improvement, the neutralization against BA.2.75.2 (98), BQ.1.1 (73), and XBB.1 (35) remain low after BA.5-bivalent-booster.

BA.5-bivalent-booster-infection sera neutralized USA-WA1/2020-, BA.4/5-, BF.7-, BA.4.6-, BA.2.75.2-, BQ.1.1-, and XBB.1-spike SARS-CoV-2s with GMTs of 5776, 1558, 1223, 744, 367, 267, and 103, respectively (**Figure 1D** and **Table S2**). The neutralizing GMTs against BA.4/5-, BF.7-, BA.4.6-, BA.2.75.2-, BQ.1.1-, and XBB.1-spike viruses were 3.7-, 4.7-, 7.8-, 15.7-, 21.6-, and 56.1-fold lower than the GMT against the USA-WA1/2020-spike SARS-CoV-2, respectively (**Figure 1D**). Compared with BA.5-bivalent-booster sera without infection history, BA.5-bivalent-booster-infection sera increased the neutralizing GMTs against USA-WA1/2020-, BA.4/5-, BF.7-, BA.4.6-, BA.2.75.2-, BQ.1.1-, and XBB.1-spike viruses by 1.6-, 5.2-, 4.0-, 4.1-, 3.7-, 3.7-, and 2.9-fold, respectively (Compare **Figure 1C** and **1D**). The results suggest that (i) previous infection significantly increases the magnitude and breadth of neutralization for BA.5-bivalent-booster and (ii) among the tested Omicron sublineages, XBB.1 exhibits the highest level of immune evasion.

Collectively, our neutralization results support three conclusions. First, the BA.5-bivalent booster elicits better neutralization against the newly emerged Omicron sublineages than the parental mRNA vaccine. Second, individuals with SARS-CoV-2 infection history develop higher and broader neutralization against the ongoing Omicron sublineages after the BA.5-bivalent booster. Third, among tested Omicron sublineages, BA.2.75.2, BQ.1.1, and XBB.1 exhibit the greatest evasion against vaccine-elicited neutralization, suggesting the potential of these new sublineages to dethrone BA.5 as the dominant lineage in circulation.

The study has two limitations. First, we have not examined the antiviral roles of nonneutralizing antibodies and cell-mediated immunity. These two immune components, together with neutralizing antibodies, protect patients from severe disease and death [16, 17]. Unlike neutralizing antibodies, many T cell epitopes after vaccination or natural infection are preserved in Omicron spikes [18]. However, robust antibody neutralization is critical to prevent viral infection [19]. Second, we have not defined the spike mutations that contribute to the observed immune evasion of the newly emerged Omicron sublineages. Spike mutation F486V was previously shown to drive the immune evasion of BA.4/5 [10]. The new Omicron sublineages BA.2.75.2, BA.4.6, BF.7, BQ.1.1, and XBB.1 share the spike R346T mutation that was reported to confer higher neutralization evasion [20]. Despite these limitations, our laboratory investigation, along with real world effectiveness of BA.5-bivalent-booster, will guide vaccine strategy against the current and future Omicron sublineages.

## Methods

### Ethical statement

All virus work was performed in a biosafety level 3 (BSL-3) laboratory with redundant fans in the biosafety cabinets at The University of Texas Medical Branch at Galveston. All personnel wore powered air-purifying respirators (Breathe Easy, 3M) with Tyvek suits, aprons, booties, and double gloves. The research protocol regarding the use of human serum specimens was reviewed and approved by the University of Texas Medical Branch (UTMB) Institutional Review Board (IRB number 20-0070). No informed consent was required since these deidentified sera were leftover specimens before being discarded. No diagnosis or treatment was involved either.

### Cells

Vero E6 (ATCC^®^ CRL-1586) purchased from the American Type Culture Collection (ATCC, Bethesda, MD) and Vero E6 cells expressing TMPRSS2 purchased from SEKISUI XenoTech, LLC were maintained in a high-glucose Dulbecco’s modified Eagle’s medium (DMEM) containing 10% fetal bovine serum (FBS; HyClone Laboratories, South Logan, UT) and 1% penicillin/streptomycin at 37°C with 5% CO_2_. Culture media and antibiotics were purchased from Thermo Fisher Scientific (Waltham, MA). The cell line was tested negative for *Mycoplasma.*

### Human Serum

Three panels of human sera were used in the study. The first panel consisted of 25 sera collected from individuals 1-3 months after receiving dose 4 of parental mRNA vaccine from Pfizer or Moderna. This panel had been tested negative for SARS-CoV-2 nucleocapsid protein expression using Bio-Plex Pro Human IgG SARS-CoV-2 N/RBD/S1/S2 4-Plex Panel (Bio-rad). The second panel consisted of 29 sera collected from individuals 1 month after BA.5-bivalent-booster of Pfizer or Moderna vaccine. All sera from this panel were tested negative for antibodies against SARS-CoV-2 nucleocapsid protein. The third panel consisted of 23 sera from individuals who were previously infected by SARS-CoV-2 (as determined by SARS-CoV-2 nucleocapsid ELISA) and received a BA.5-bivalent-booster 1 month before serum collection. The genotypes of the infecting SARS-CoV-2 variants could not be determined for the third serum panel. Patient information was completely deidentified from all specimens. The deidentified human sera were heat-inactivated at 56°C for 30 min before the neutralization test. The serum information is presented in Tables S1-3.

### Generation of recombinant Omicron sublineages-mNG SARS CoV-2 viruses

Recombinant Omicron sublineage BA.4/5-, BF.7-, BA.4.6-, BA.2.75.2-, BQ.1.1-, and XBB.1-spike mNG SARS-CoV-2s was constructed by engineering the complete spike gene from the indicated variants into an infectious cDNA clone of mNG USA-WA1/2020 as reported previously [10, 13]. Spike sequences were based on BA.4/5 (BA.4: GISAID EPI_ISL_11 542270; BA.5: GISAID EPI_ISL_11542604; BA.4 and BA.5 have the identical spike sequence), BA.4.6 (GISAID EPI_ISL_15380489), BA.2.75.2 (GISAID EPI_ISL_14458978), BF.7 (GISAID EPI_ISL_14425795), BQ.1.1 (GISAID EPI_ISL_15542649) and XBB.1 (GISAID EPI_ISL_15232105). The full-length infectious cDNA clone of SARS-CoV-2 was assembled by *in vitro* ligation followed by *in vitro* transcription to synthesize the viral genomic RNA. The fulllength RNA transcripts were electroporated in Vero E6-TMPRSS2 cells to recover the viruses. Viruses were rescued post 2-3 days after electroporation and served as P0 stock. P0 stock was further passaged once on Vero E6 cells to produce P1 stock. The spike gene was sequenced from all P1 stock viruses to ensure no undesired mutation. The infectious titer of the P1 virus was quantified by fluorescent focus assay on Vero E6 cells. The P1 virus was used for the neutralization test. The protocols for the mutagenesis of mNG SARS-CoV-2 and virus production were reported previously [21]. All virus preparation and infections were carried out at the biosafety level 3 (BSL-3) facility at the University of Texas Medical Branch at Galveston.

### Fluorescent focus reduction neutralization test (FFRNT)

Neutralization titers of human sera were measured by FFRNT using the USA-WA1/2020-, BA.4/5-, BF.7-, BA.4.6-, BA.2.75.2-, BQ.1.1-and XBB.1-spike mNG SARS-CoV-2s. The details of the FFRNT protocol were reported previously [3]. Briefly, 2.5 × 10^4^ Vero E6 cells per well were seeded in 96-well plates (Greiner Bio-one™). The cells were incubated overnight. On the next day, each serum was 2-fold serially diluted in the culture medium with the first dilution of 1:20 (final dilution range of 1:20 to 1:20,480). The diluted serum was incubated with 100-150 FFUs of mNG SARS-CoV-2 at 37 °C for 1 h, after which the serum virus mixtures were loaded onto the pre-seeded Vero E6 cell monolayer in 96-well plates. After 1 h infection, the inoculum was removed and 100 μl of overlay medium (supplemented with 0.8% methylcellulose) was added to each well. After incubating the plates at 37 °C for 16 h, raw images of mNG foci were acquired using CytationTM 7 (BioTek) armed with 2.5× FL Zeiss objective with a wide-field of view and processed using the Gene 5 software settings (GFP [469,525] threshold 4000, object selection size 50-1000 μm). The foci in each well were counted and normalized to the nonserum-treated controls to calculate the relative infectivities. The FFRNT_50_ value was defined as the minimal serum dilution that suppressed >50% of fluorescent foci. The neutralization titer of each serum was determined in duplicate assays, and the geometric mean was taken. All attempts at replication were successful. Tables S1-3 summarize the FFRNT50 results. Data were initially plotted in GraphPad Prism 9 software and assembled in Adobe Illustrator.

### Data availability

The raw data that support the findings of this study are shown in the Source data files.

## Acknowledgments

We thank colleagues at the University of Texas Medical Branch (UTMB) for helpful discussions P.-Y.S. was supported by NIH contract HHSN272201600013C, and awards from the Sealy & Smith Foundation, the Kleberg Foundation, the John S. Dunn Foundation, the Amon G. Carter Foundation, the Summerfield Robert Foundation, and Edith and Robert Zinn.

## Author contributions

Conceptualization, P.R., X.X., P.-Y.S.; Methodology, C.K., J.Z., H.X., M.L., H.C.C., P.R., X.X., P.-Y.S.; Investigation, C.K., J.Z., H.X., M.L., H.C.C., P.R., X.X., P.-Y.S.; Resources, P.R., X.X., P.-Y.S.; Data Curation, C.K., J.Z., P.R., X.X.; Writing-Original Draft, P.R., X.X., P.-Y.S.; Writing-Review & Editing, C.K., J.Z., H.X., M.L., H.C.C., P.R., X.X., P.-Y.S.; Supervision, P.R., X.X., P.-Y.S.; Funding Acquisition, P.R., X.X., P.-Y.S.

