## Supplementary Tables 1-3 for "Low neutralization of SARS-CoV-2 Omicron BA.2.75.2, BQ.1.1, and XBB.1 by 4 doses of parental mRNA vaccine or a BA.5-bivalent booster" (from another version of this preprint: version 1, 2022-11-02)

**Table S1.** Twenty-five human serum samples collected after dose 4 of parental mRNA vaccine.

| Serum ID | Age (year) | Gender (F/M) | Ethnicity | Serum collection time (days post-dose 4 vaccination) | mRNA Vaccine type | *FFRNT_50_ | | | | | | |
| --- | --- | --- | --- | --- | --- | --- | --- | --- | --- | --- | --- | --- |
|  |  |  |  |  |  | USA-WA1/2020 | BA.4/5-spike | BF.7-spike | BA.4.6-spike | BA.2.75.2-spike | BQ.1.1-spike | XBB.1-spike |
| 1 | 62 | F | White | 37 | Pfizer | 320 | ^10 | 10 | 10 | 10 | 10 | 10 |
| 2 | 80 | M | Hispanic | 30 | Pfizer | 10240 | 320 | 80 | 80 | 40 | 20 | 20 |
| 3 | 84 | F | White | 55 | Pfizer | 1280 | 80 | 14 | 10 | 10 | 10 | 10 |
| 4 | 92 | F | White | 27 | Pfizer | 2560 | 113 | 40 | 40 | 28 | 10 | 10 |
| 5 | 78 | M | White | 56 | Pfizer | 320 | 10 | 10 | 14 | 10 | 10 | 10 |
| 6 | 71 | F | White | 27 | Moderna (dose 1-3), Pfizer (dose 4) | 2560 | 320 | 320 | 320 | 80 | 80 | 40 |
| 7 | 83 | F | White | 23 | Pfizer | 1280 | 160 | 113 | 80 | 20 | 20 | 10 |
| 8 | 87 | M | White | 24 | Pfizer | 1280 | 113 | 226 | 160 | 10 | 57 | 10 |
| 9 | 80 | M | White | 52 | Pfizer | 5120 | 640 | 320 | 226 | 80 | 80 | 40 |
| 10 | 84 | M | White | 26 | Pfizer | 640 | 20 | 10 | 10 | 10 | 10 | 10 |
| 11 | 75 | M | Black | 47 | Pfizer | 1810 | 57 | 40 | 40 | 226 | 20 | 40 |
| 12 | 90 | M | Black | 34 | Pfizer | 2560 | 160 | 80 | 80 | 40 | 40 | 20 |
| 13 | 59 | F | Hispanic | 27 | Pfizer (dose 1-3), Moderna (dose 4) | 1810 | 160 | 113 | 160 | 10 | 28 | 10 |
| 14 | 72 | F | White | 52 | Pfizer | 1810 | 40 | 40 | 40 | 20 | 14 | 10 |
| 15 | 73 | M | White | 94 | Moderna | 640 | 40 | 40 | 28 | 10 | 10 | 10 |
| 16 | 67 | F | Black | 50 | Pfizer | 1280 | 80 | 80 | 80 | 20 | 40 | 10 |
| 17 | 75 | F | White | 78 | Pfizer | 2560 | 453 | 320 | 320 | 80 | 80 | 40 |
| 18 | 86 | M | White | 49 | Pfizer | 1810 | 320 | 226 | 320 | 40 | 20 | 20 |
| 19 | 66 | F | White | 48 | Pfizer | 453 | 20 | 20 | 14 | 10 | 10 | 10 |
| 20 | 80 | M | Black | 44 | Pfizer | 1280 | 160 | 160 | 160 | 40 | 28 | 20 |
| 21 | 78 | M | White | 73 | Pfizer | 905 | 40 | 40 | 28 | 20 | 10 | 10 |
| 22 | 86 | M | White | 51 | Pfizer | 1280 | 20 | 20 | 20 | 10 | 10 | 10 |
| 23 | 84 | F | White | 35 | Pfizer | 20480 | 1280 | 640 | 453 | 226 | 160 | 80 |
| 24 | 94 | F | White | 47 | Pfizer | 1280 | 80 | 40 | 40 | 20 | 20 | 10 |
| 25 | 87 | F | Hispanic | 43 | Pfizer | 905 | 160 | 320 | 160 | 28 | 14 | 10 |
| ^#^GMT | - | - | - | - | - | 1533 | 95 | 69 | 62 | 26 | 22 | 15 |
| ^†^95% CI | - | - | - | - | - | 1036-2268 | 56-160 | 41-115 | 38-103 | 17-38 | 16-31 | 12-20 |

*Individual FFRNT_50_ value is the geometric mean of duplicate FFRNT results.

^FFRNT_50_ of <20 was treated as 10 for plot purpose and statistical analysis.

^#^Geometric mean neutralizing titers (GMT).

^†^95% confidence interval (95% CI) for the GMT.

**Table S2.** Twenty-nine human serum samples collected one month after BA.5-bivalent-booster without SARS-CoV-2 infection history.

| Serum ID | Age (year) | Gender (F/M) | Ethnicity | Serum collection time (days post-BA.5-bivalent booster) | Last dose of mRNA vaccine before BA.5-bivalent booster | *FFRNT_50_ | | | | | | |
| --- | --- | --- | --- | --- | --- | --- | --- | --- | --- | --- | --- | --- |
|  |  |  |  |  |  | USA-WA1/2020 | BA.4/5-spike | BF.7-spike | BA.4.6-spike | BA.2.75.2-spike | BQ.1.1-spike | XBB.1-spike |
| 1 | 34 | F | Black | 15 | Dose 3 | 5120 | 640 | 1280 | 640 | 640 | 226 | 160 |
| 2 | 78 | F | White | 17 | Dose 4 | 5120 | 640 | 640 | 320 | 320 | 160 | 80 |
| 3 | 86 | M | White | 21 | Dose 4 | 1280 | 28 | 10 | 10 | 20 | 10 | 10 |
| 4 | 31 | M | Asian | 20 | Dose 2 | 5120 | 1280 | 1280 | 905 | 160 | 160 | 80 |
| 5 | 61 | F | White | 15 | Dose 4 | 5120 | 320 | 453 | 320 | 160 | 80 | 57 |
| 6 | 58 | F | Black | 14 | Dose 3 | 7241 | 1810 | 1810 | 1280 | 320 | 640 | 113 |
| 7 | 69 | M | White | 22 | Dose 4 | 2560 | 80 | 80 | 57 | 57 | 20 | 14 |
| 8 | 67 | F | Asian | 21 | Dose 4 | 20480 | 2560 | 1280 | 640 | 160 | 160 | 40 |
| 9 | 77 | M | Asian | 27 | Dose 4 | 2560 | 226 | 320 | 160 | 80 | 40 | 40 |
| 10 | 39 | F | White | 15 | Dose 3 | 5120 | 320 | 320 | 226 | 80 | 160 | 20 |
| 11 | 73 | M | White | 24 | Dose 4 | 1810 | 160 | 160 | 80 | 80 | 40 | 28 |
| 12 | 83 | F | White | 17 | Dose 4 | 1280 | 40 | 80 | 40 | 40 | 10 | 10 |
| 13 | 79 | F | White | 26 | Dose 3 | 3620 | 320 | 640 | 320 | 226 | 57 | 57 |
| 14 | 35 | F | Asian | 29 | Dose 3 | 5120 | 160 | 320 | 226 | 80 | 80 | 40 |
| 15 | 73 | M | White | 26 | Dose 3 | 7241 | 320 | 320 | 160 | 80 | 80 | 28 |
| 16 | 76 | M | White | 32 | Dose 4 | 7241 | 1280 | 905 | 640 | 226 | 160 | 160 |
| 17 | 71 | M | White | 28 | Dose 4 | 1280 | 40 | 57 | 40 | 40 | 20 | 10 |
| 18 | 22 | M | Hispanic | 19 | Dose 3 | 10240 | 2560 | 1280 | 1280 | 320 | 453 | 80 |
| 19 | 61 | F | White | 30 | Dose 4 | 640 | 80 | 80 | 57 | 20 | 28 | 14 |
| 20 | 56 | M | White | 21 | Dose 3 | 10240 | 640 | 640 | 453 | 160 | 160 | 80 |
| 21 | 66 | F | White | 26 | Dose 4 | 3620 | 320 | 320 | 160 | 80 | 80 | 40 |
| 22 | 76 | F | White | 30 | Dose 4 | 3620 | 226 | 226 | 160 | 10 | 80 | 10 |
| 23 | 61 | F | Asian | 31 | Dose 3 | 10240 | 5120 | 5120 | 2560 | 640 | 453 | 226 |
| 24 | 77 | F | Black | 31 | Dose 4 | 5120 | 80 | 80 | 40 | 40 | 28 | 10 |
| 25 | 59 | F | unknown | 28 | Dose 4 | 2560 | 320 | 320 | 160 | 160 | 80 | 28 |
| 26 | 71 | M | Hispanic | 22 | Dose 4 | 1280 | 160 | 160 | 80 | 80 | 80 | 14 |
| 27 | 70 | F | White | 22 | Dose 3 | 1280 | 113 | 57 | 28 | 20 | 20 | 10 |
| 28 | 79 | F | White | 25 | Dose 4 | 2560 | 226 | 160 | 80 | 57 | 10 | 20 |
| 29 | 79 | M | White | 18 | Dose 4 | 2560 | 160 | 320 | 160 | 226 | 160 | 80 |
| ^#^GMT | - | - | - | - | - | 3620 | 298 | 305 | 183 | 98 | 73 | 35 |
| ^†^95% CI | - | - | - | - | - | 2668-4912 | 181-490 | 185-503 | 111-299 | 66-146 | 47-112 | 24-50 |

*Individual FFRNT_50_ value is the geometric mean of duplicate FFRNT results.

^#^Geometric mean neutralizing titers (GMT).

^†^95% confidence interval (95% CI) for the GMT.

**Table S3.** Twenty-three human serum samples collected one month after BA.5-bivalent-booster with SARS-CoV-2 infection history.

| Serum ID | Age (year) | Gender (F/M) | Ethnicity | Serum collection time (days post-BA.5-bivalent booster) | Last dose of mRNA vaccine before BA.5-bivalent booster | *FFRNT_50_ | | | | | | |
| --- | --- | --- | --- | --- | --- | --- | --- | --- | --- | --- | --- | --- |
|  |  |  |  |  |  | USA-WA1/2020 | BA.4/5-spike | BF.7-spike | BA.4.6-spike | BA.2.75.2-spike | BQ.1.1-spike | XBB.1-spike |
| 1 | 19 | F | White | 19 | Dose 3 | 10240 | 1280 | 1280 | 640 | 640 | 160 | 113 |
| 2 | 69 | F | White | 15 | Dose 4 | 10240 | 5120 | 5120 | 2560 | 905 | 1280 | 320 |
| 3 | 59 | F | White | 15 | Dose 3 | 20480 | 10240 | 10240 | 7241 | 2560 | 2560 | 453 |
| 4 | 46 | F | White | 18 | Dose 2 | 5120 | 2560 | 2560 | 640 | 320 | 320 | 160 |
| 5 | 80 | F | Black | 22 | Dose 3 | 10240 | 7241 | 5120 | 3620 | 2560 | 640 | 320 |
| 6 | 76 | F | Hispanic | 21 | Dose 3 | 5120 | 1280 | 1280 | 905 | 453 | 160 | 113 |
| 7 | 67 | M | White | 15 | Dose 4 | 5120 | 2560 | 1280 | 1280 | 320 | 320 | 80 |
| 8 | 52 | F | White | 30 | Dose 4 | 5120 | 1280 | 1280 | 905 | 160 | 640 | 160 |
| 9 | 48 | F | Asian | 24 | Dose 3 | 14482 | 7241 | 5120 | 2560 | 1280 | 640 | 320 |
| 10 | 58 | F | White | 28 | Dose 2 | 5120 | 640 | 640 | 320 | 160 | 226 | 40 |
| 11 | 67 | F | Black | 25 | Dose 2 | 5120 | 2560 | 2560 | 1280 | 640 | 640 | 160 |
| 12 | 20 | F | Hispanic | 21 | Dose 2 | 1810 | 320 | 453 | 160 | 80 | 160 | 28 |
| 13 | 75 | M | White | 21 | Dose 3 | 20480 | 2560 | 2560 | 1280 | 1280 | 453 | 320 |
| 14 | 64 | F | White | 19 | Dose 4 | 7241 | 2560 | 2560 | 1280 | 2560 | 453 | 320 |
| 15 | 90 | F | White | 22 | Dose 3 | 10240 | 1280 | 1280 | 1280 | 320 | 640 | 160 |
| 16 | 35 | F | Hispanic | 32 | Dose 2 | 1280 | 320 | 226 | 160 | 80 | 80 | 28 |
| 17 | 39 | M | Asian | 31 | Dose 3 | 10240 | 5120 | 2560 | 2560 | 640 | 320 | 80 |
| 18 | 67 | M | White | 17 | Dose 4 | 10240 | 2560 | 1810 | 1280 | 640 | 640 | 226 |
| 19 | 68 | M | Hispanic | 14 | Dose 3 | 5120 | 2560 | 1280 | 905 | 320 | 160 | 80 |
| 20 | 51 | M | White | 28 | Dose 3 | 7241 | 905 | 320 | 320 | 320 | 160 | 57 |
| 21 | 65 | M | Asian | 29 | Dose 4 | 2560 | 453 | 160 | 80 | 113 | 57 | 40 |
| 22 | 64 | M | White | 16 | Dose 3 | 2560 | 640 | 640 | 320 | 40 | 40 | 20 |
| 23 | 73 | M | Black | 31 | Dose 4 | 640 | 57 | 40 | 28 | 20 | 10 | 10 |
| ^#^GMT | - | - | - | - | - | 5776 | 1558 | 1223 | 744 | 367 | 267 | 103 |
| ^†^95% CI | - | - | - | - | - | 3994-8352 | 922-2631 | 704-2125 | 425-1301 | 209-644 | 158-452 | 66-162 |

*Individual FFRNT_50_ value is the geometric mean of duplicate FFRNT results.

^#^Geometric mean neutralizing titers (GMT).

^†^95% confidence interval (95% CI) for the GMT.
